# Identification of age and ethnicity specific gene expression biomarkers for immune aging

**DOI:** 10.1101/2021.02.22.432179

**Authors:** Yang Hu, Yudai Xu, Lipeng Mao, Wen Lei, Jian Xiang, Guodong Zhu, Yutian Hu, Haitao Niu, Feng Gao, Lijuan Gao, Li’an Huang, Oscar Junhong Luo, Guobing Chen

## Abstract

Human immune system functions over an entire lifetime, yet how and why the immune system becomes less effective with age are not well understood. Here, we characterize peripheral blood mononuclear cells transcriptome from 172 healthy adults with 21~90 years of age using RNA-seq and the weighted gene correlation network analyses (WGCNA). These data reveal a set of insightful gene expression modules and representative gene biomarkers for human immune system aging from Asian and Caucasian ancestry, respectively. Among them, the aging-specific modules show an age-related gene expression variation spike around early-seventies. In addition, it is not known whether Asian and Caucasian immune systems go through similar gene expression changes throughout their lifespan, and to what extent these aging-associated changes are shared among ethnicities. We find the top hub genes including NUDT7, CLPB, OXNAD1 and MLLT3 are shared between Asian and Caucasian aging related modules and further validated in human PBMCs from different age groups. Overall, the impact of age and race on <u>transcriptional variation</u> elucidated from this study provide insights into the transcriptional driver of immune aging.

## INTRODUCTION

The world is witnessing a rapid demographic shift towards an older population, a trend with major medical, social, economic, and political implications. Aging is a multifaceted process, involving numerous molecular and cellular mechanisms in the context of different organ systems [1]. A crucial component of aging is a set of functional and structural alterations in the immune system that can manifest as a decreased ability to fight infection, diminished response to vaccination, increased incidence of cancer, higher prevalence of autoimmunity and constitutive low-grade inflammation, among others [2]. In addition to alterations in the stromal microenvironment in primary and secondary lymphoid organs, cell-intrinsic changes in both innate and adaptive immune cells play an important role in age-associated immune dysfunction. These alterations and changes manifest themselves in increased morbidity and mortality of older organisms due to infectious disease. However, the interplay between the PBMC age-related gene expression changes that affect the immune aging remains incompletely elucidated, and there is no clear understanding of which gene changes are primary, arising as a consequence of aging, and which might be secondary, adaptive or compensatory to the primary changes. Thus, a transcriptome analyses may lend greater insight than a static genetic investigation. In contrast to the relatively invariable genome sequence, the transcriptome is highly dynamic and changes in response to stimuli. Therefore, the aim of this study was to exploit a large-scale population-based strategy to systematically identify genes and pathways differentially expressed as a function of chronological age.

More importantly, analyses of human blood samples from different race and ethnicity uncovered significant aging-related variations in various subsets of PBMCs [3]. For example, a study on PBMC subsets characterized not only the presence of benign ethnic neutropenia among African Americans but further discovered a higher proportion of CD 19+ cells and a lower proportion of CD3+ cells than in Caucasian population [4]. Moreover, the proportions of PBMCs’ subpopulation in Asian cohorts were also different. Choong and colleagues observed that there were differences in cell counts for T, NK, and CD4+ cells as well as in the CD4/CD8 ratio among healthy Malaysians, Chinese, and Indians across the life span (18~71 years) [5]. In addition, Indians were significantly different from Malays and Chinese. Indians had higher T cells, higher CD4 cells, higher CD4/CD8 ratio, and lower NK cells; Chinese donors had lower B-cell levels than Malays and Indians [5]. Despite the importance of age and race in shaping immune cell numbers and functions, it is not known whether Asian and Caucasian immune systems go through similar gene expression changes throughout their lifespan, and to what extent these aging-associated changes are shared among ethnicities.

To study this, we compared the PBMC transcriptomes of healthy Asian and Caucasian adults with matching aging-stage by RNA-Seq. Computational WGCNA analyses, differentially expressed genes (DEGs) and functional enrichments analyses revealed distinct immune system aging signatures in Asian and Caucasian. These findings uncovered in which ways aging differentially affects the immune systems between ethnicities and discovered a common genetic variant that greatly impacts normal PBMC aging.

## RESULTS

### Transcriptome profiling of PBMCs from donors with distinct ancestry

We recruited 19 community dwelling healthy volunteers (10 female, 9 male) whose ages span 21~93 years old (Supplementary Table 1): 9 young (age 21~30: 5 female, 9 male), and 10 old donors (age ≥74: 5 female, 5 male), from Guangdong Province of China. No significant differences were detected between sexes in their frailty scores or age distributions. Male and female samples for each assay were comparable in terms of age (males: ~81.4 vs. females: ~85.2; t-test *p*-value = 0.41) (Supplementary Table 1). Then, PBMCs transcriptome of these 19 donors were profiled using RNA-seq. As a result, 29,367 genes were selected after normalization of raw expression counts, excluding the genes with no expression in all samples. Moreover, transcriptome data of a total of 153 normal healthy human subjects, whose ages span 20~90 years old from different ethnicities (Figure S1, Supplementary Table 2): 65 young (ages 21~40: 24 males, 41 females), 40 middle-aged (ages 41~64: 18 men, 22 women), and 48 older subjects (65+: 20 men, 28 women) were downloaded from The 10,000 Immunomes Project (10KIP, http://10kimmunomes.org/). Then, their PBMCs gene expression data were processed for further WGCNA clustering analyses.

### Aging and race cause transcriptomic variations over human adult lifespan

To identify major sources of variation in transcriptomic data, we conducted WGCNA analyses using expressed genes (n = 19,089) from the 10KIP. First, by using the soft thresholding power (β=6) in this algorithm, the co-expression network could satisfy the approximate scale-free topology criterion with R^2^ > 0.80 while maintaining a high mean connectivity with enough information (Figure S2A). Then, we merged the modules of eigengenes with a correlation coefficient of over 0.75 (Figure S2B), and the size of the co-expression modules ranged from 68 to 3,891 genes (Figure 1A). After merging the highly correlated modules, the co-expression genes were clustered into 22 modules and labeled with different colors (Figure 1B). To further quantify co-expression similarity of all modules, we calculated their eigengenes adjacency on their correlation of the entire modules. Each module showed independent validation to each other, and higher correlation indicated higher co-expression interconnectedness (Figure 1B). Genes within the same module exhibited higher correlation than the genes between different modules.

**Figure 1.**
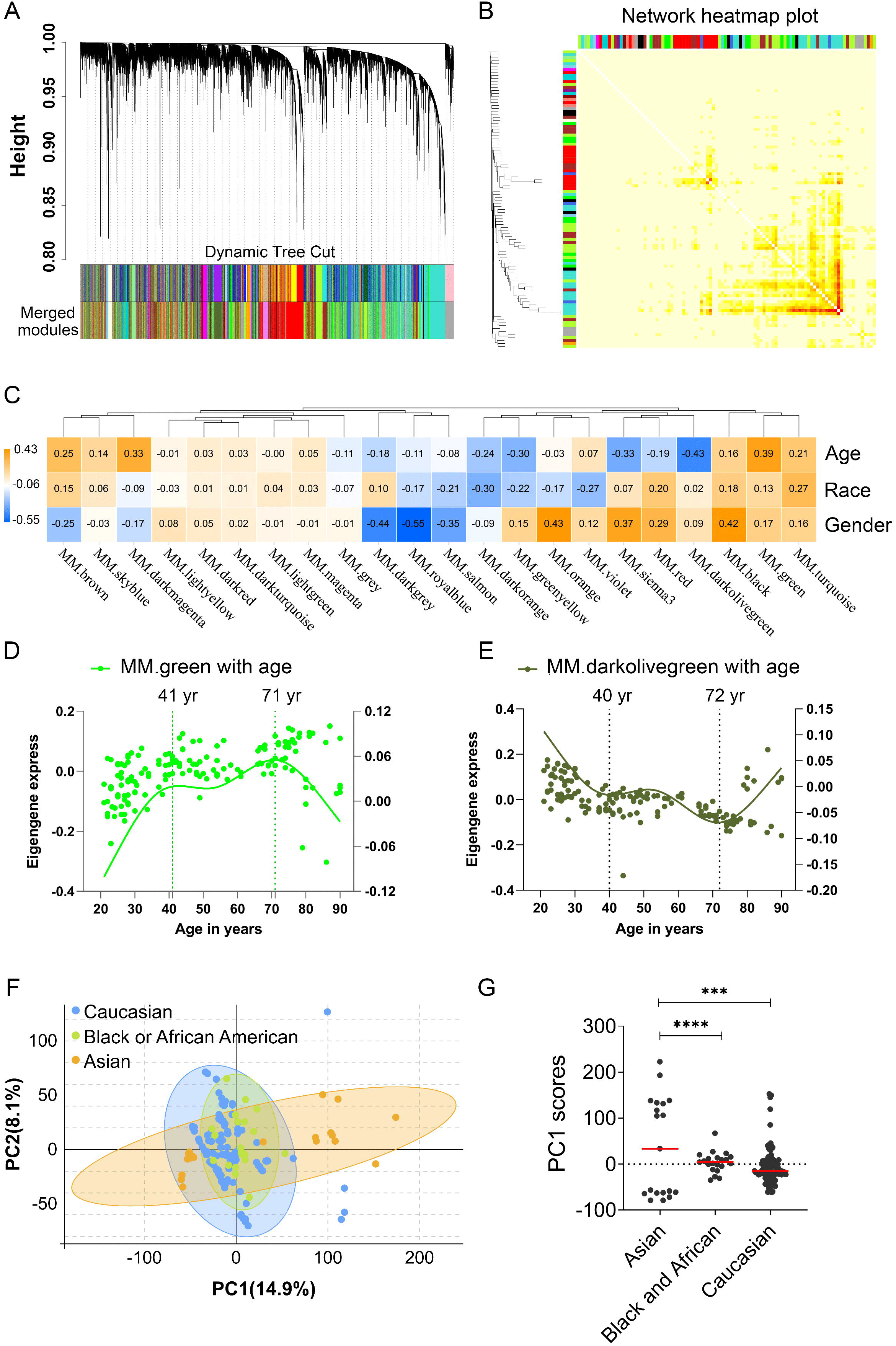
Age and race influenced the transcriptomic variations over human adult lifespan. WGCNA approach was applied for gene module consturction for the transcriptome data of 153 healthy human subjects in 10KIP. Principal component analyses (PCA) were calculated individually. (A) Cluster dendrogram. Each branch represented one gene and each color below denoted one co-expression gene module. The two colored rows below the dendrogram represented the original and merged modules, respectively. (B) Eigengene adjacency heatmap of different modules. Each module showed independent validation to each other, and higher correlation indicated higher co-expression interconnectedness. (C) Heatmap of the correlation between trait (age, race and gender) and module eigengenes (ME, n=22). The column and row corresponded to ME and trait, respectively. Each cell contained the value of Pearson’s correlation coefficient. The table was color-coded by correlation according to the color legend. The *p*-value < 0.05 represented statistical significance. (D-E) The characteristic gene expression changed during PBMC aging. The left and right-hand Y-axis represented the eigengene expression of each module, and trend line for each individuals, respectively. (F-G) Principal component1 scores (PC1) were calculated for each individual from principal component analyses (PCA). PC1 scores from transcriptomic data were differentially expressed among different races.

In order to identify which factor mainly affected the variation in PBMC transcriptomes, the correlation coefficients between modules and the trait of age, gender and race were calculated, respectively. These traits moderately correlated with one or more modules, and the trait-module relationships with P ≤ 0.01 were further analyzed (Figure 1C, Supplementary Table 3). Notably, among them, green module significantly correlated with age positively (Pearson r=0.39, *p*=8.30e-07; Figure 1C), while darkolivegreen module negatively correlated with age (Pearson r=0.45, *p*=2.02e-08; Figure 1C). Furthermore, a significant correlation was also detected between sex and race in terms of their transcriptomic aging signatures respectively. To study whether PBMC transcriptomic changes were acquired gradually over lifespan or more rapidly at certain ages, we detected age brackets during which abrupt changes take place, referred as breakpoints. Within each aging related module, we compared transcriptomic profiles observed at different ages (n=153). Finally, these analyses revealed two periods in adult lifespan during which rapid changes occurred: (i) a timepoint in early-forties, and (ii) a later timepoint after 70 years of age (Figure 1D, E, Supplementary Table 4).

Besides, racial/ethnic differences in PBMC aging among adult lifespan were also important with profound effect on health. To identify major sources of variation in transcriptomic data, we conducted the principal component analyses (PCA) using expressed genes (n = 19,089) from high-quality samples. The first principal component (PC1) captured 14.9% of the variation in 153 microarray data and associated to age groups (Figure 1F, Supplementary Table 5). The variants of PC1 score between Asian and Caucasian samples were more significantly different (Figure 1G), and this also took place in Asian and Black or Africa American (Figure 1G). Together, these results suggested that aging and race had both influenced the variation in PBMC transcriptomes, where it was unclear to what extent these aging-associated changes were shared in different races, such as Asian and Caucasian.

### Aging-related PBMC transcriptome dynamics in Asian (Chinese)

PCA of the 19 PBMC transcriptomes of Asian donors (Chinese) revealed that young and old samples were divided into two parts, and females had larger variants than males, especially in the old group (Figure 2A). To further identify the related genes of PBMC aging, WGCNA was conducted using FPKM values of 29,367genes (FPKM > 1 of all sequenced transcript) and the trait of age and sex (Figure 2B). Genes with the same expression pattern were clustered into the same module to generate a cluster dendrogram (Figure 2B). The sample dendrogram and trait heatmap were visualized to understand the relationship between the corresponding gene expression data and biological traits (Figure 2C). Forty modules were obtained, of which four ME-modules (cyan, darkturquoise, orange, brown) showed significant correlations with age, with the absolute Pearson correlation coefficients |R| ≥ 0.70 ( *p* < 0.01) (Figure 2C, Supplementary Table 6). Further, a consensus clustering also confirmed the four main modules were clearly separated between young to old (Figure 2D). Similarly, based on the module eigengene (ME) expression profile and the ages of the donors, these four significant modules all showed sharp changes at the age of 74 (Figure 2E, Supplementary Table 7). These results suggested these four gene modules were highly associated with chronological age in Asian (Chinese), especially for the brown (r=0.85, *p*=3.54E-06) and darkturquoise modules (r=0.77, *p*=9.90E-05).

**Figure 2.**
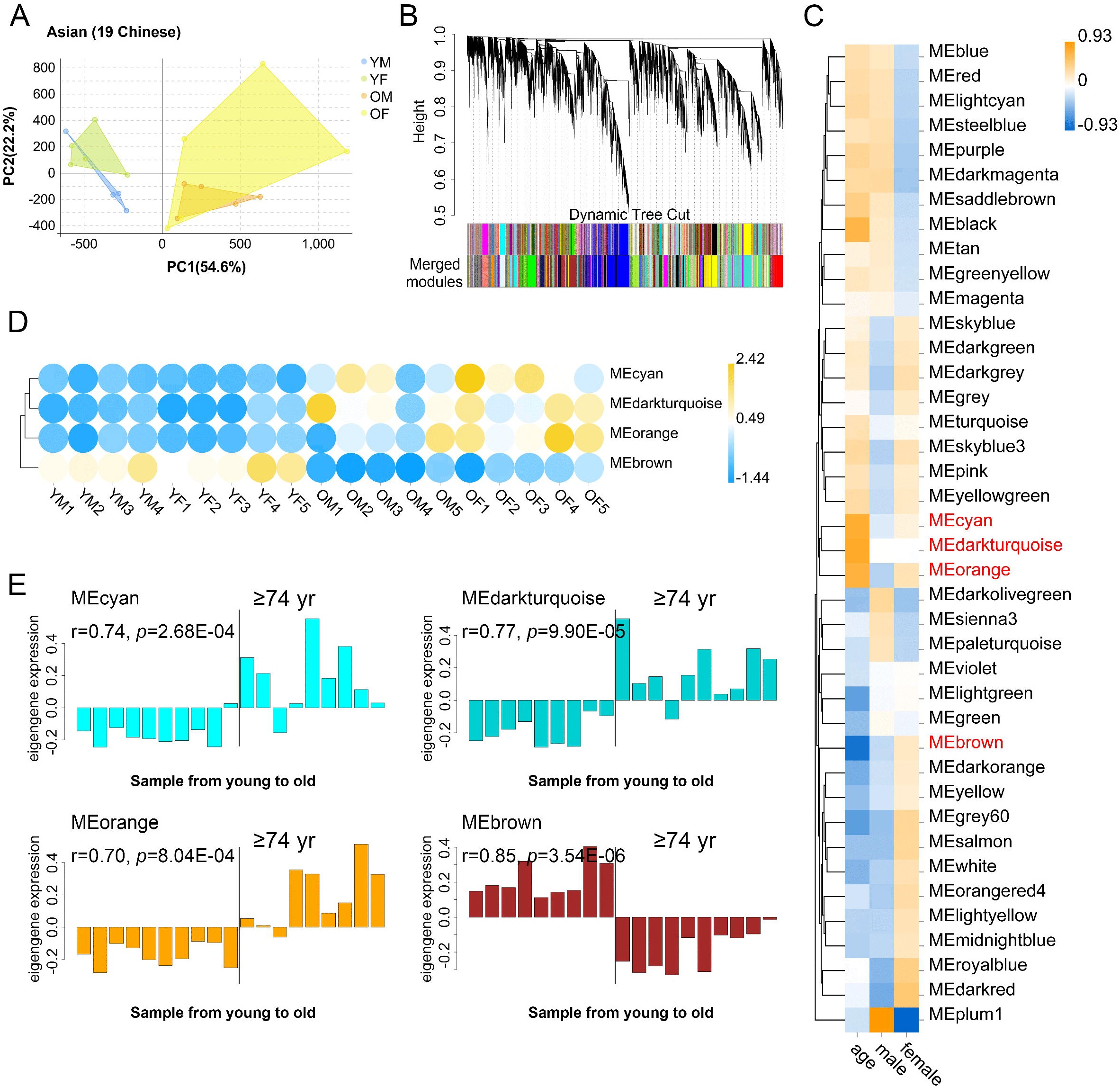
The characteristic gene expression of PBMC aging in Asian (Chinese). Transcriptome data of 19 healthy human subjects in Guangdong China were analyzed, and gene modules were constructed by WGCNA. (A) Principal component analyses. Young and old individuals were largely separated according to the principal component1 scores (PC1). (B) Cluster dendrogram. Each branch represents one gene and each color below denotes one co-expression gene module. The two colored rows below the dendrogram, represented the original and merged modules, respectively; (C) Heatmap of the correlation between trait (age and gender) and module eigengenes (ME, n=40). The column and row corresponded to trait and ME, respectively. The color in each cell represented corresponding correlation, and scaled in the color legend. (D) Hierarchical cluster analyses of four interested modules. Based on the module-trait’s correlation and *p* value (absolute r > 0.5, *p*< 0.05), three modules (cyan, darkturquoise, and orange) showed relatively lower expression in young adults and high expression in the aged population. Conversely, the brown modules showed the opposite result. Each circle represented an individual. (E) The histograms of the eigengene expression in the four age-related modules from young to old.

### Novel and known age-associated genes and pathways associated with PBMC aging in Asian (Chinese)

WGCNA analyses defined that the module eigengene (ME) was the first principal component of a given module and could be considered as a representative of the module’s gene expression profile. Based on ME expression profile of the four significant modules, the expression of cyan, darkturquoise and orange modules were downregulated in young donors, while brown module showed the opposite results (Figure 3A). To further explore the biological functions of the most closely age-related modules (brown module r=0.85, *p*=3.54E-06; darkturquoise module r=0.77, *p*=9.90E-05), we performed Gene Ontology (GO) enrichment analyses, as well as pathway ontology analyses by using clusterProfiler R package [6] (Supplementary Table 8). The enrichment analyses revealed that in the brown module, the top two enriched terms in GO ontology were “Cellular amino acid metabolic process” (FDR=5.74E-04) and “Negative regulation of neuron apoptotic process” (FDR=8.43E-04) (Figure 3B); for the KEGG pathway analyses, the top enriched terms were “Herpes simplex virus 1 infection” (FDR= 9.19E-09) and “Valine, leucine and isoleucine degradation” (FDR=1.33E-03) (Figure 3C). Meanwhile, functional annotations of darkturquoise module genes showed the top enriched terms in the GO databases were “Protein-DNA complex subunit organization” (FDR=7.68E-07) and “ncRNA processing” (FDR=1.33E-06) (Figure 3B). Moreover, genes in darkturquoise module were found to be significantly enriched in protein export and lysine degradation signaling pathway (Figure 3C). These findings together with previous research, which found persistent virus infections and metabolic dysregulation were closely related with immune aging [7], implied that the above signaling pathways might play an important role in aging.

**Figure 3.**
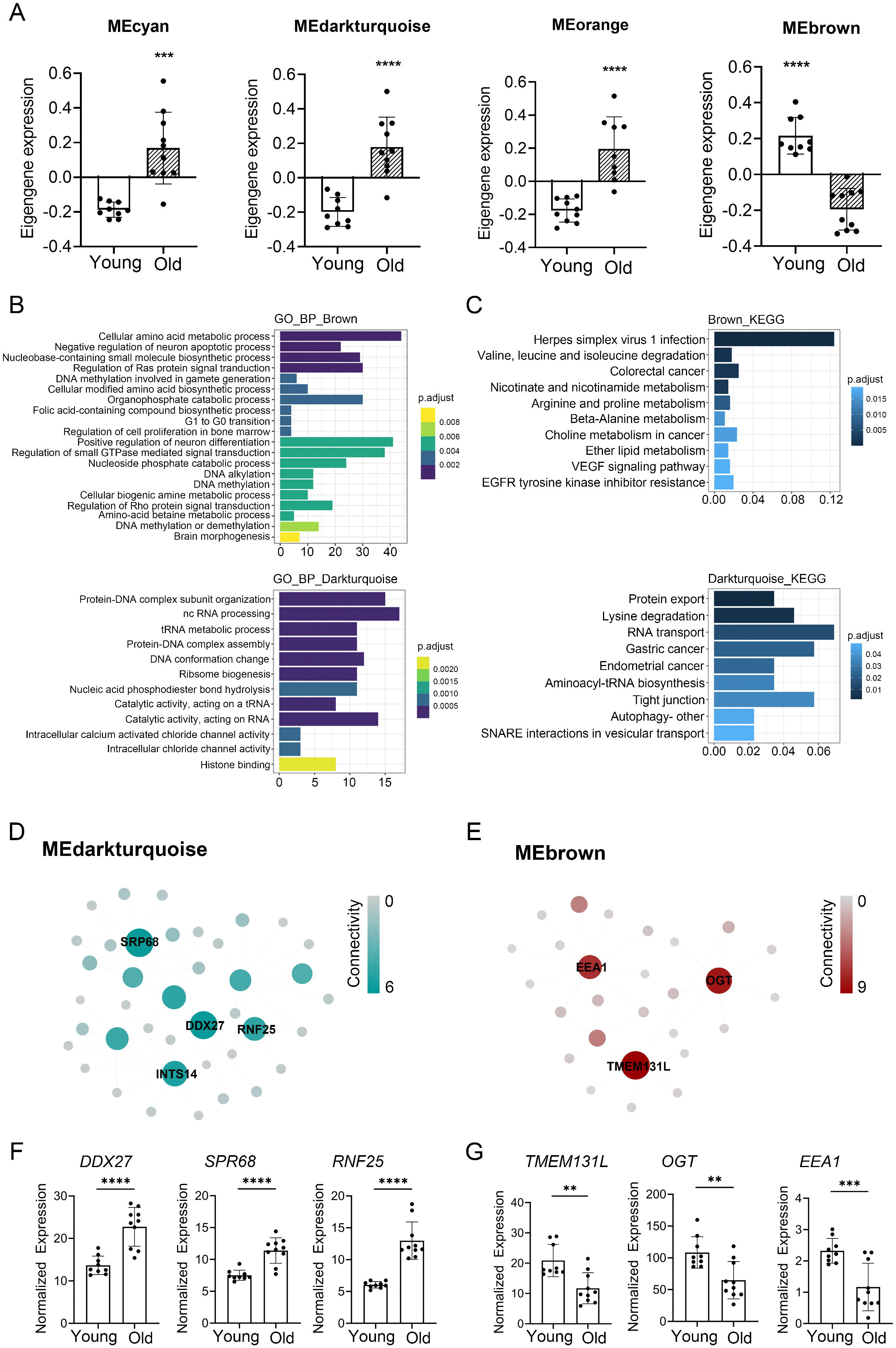
Novel and known age-associated genes and pathways associated with PBMC aging in Asian (Chinese). Gene Ontology (GO) and KEGG pathway enrichment analyses were conducted to analyze the biological functions of modules. Module eigengene (ME) was the first principal component of a given module and could be considered as a representative of the module’s gene expression profile. (A) The transcriptomic expression of age related modules changed significantly among young and old individuals. Based on ME expression profile of the four interesting modules, the expression of cyan, darkturquoise and orange modules were downregulated, while brown module showed the opposite results. The *p*-value was calculated by the student’s t-test, n=19, \**p* < 0.05, \*\**p* < 0.01, \*\*\**p* < 0.001, \*\*\*\**p* < 0.0001. (B-C) Top 10 GO biological process functional annotation (B) and KEGG pathway enrichment (C) analyses for the brown and darkturquoise modules. The color represented the adjusted *p*-values. (D-E) Hub gene detection for the darkturquoise (D) and brown (E)modules. PPI network of the brown and darkturquoise modules were based on the STRING database. And each node represented a protein-coding gene and the size of each node was mapped to its connectivity (also known as degree). (F-G) The verification of the hub genes. The top of three genes in brown and darkturquoise modules were selected and its mRNA abundance of these hub genes were detected in young and old individuals. The *p*-value was calculated by the student’s t-test, young=9, old=10, \**p* < 0.05, \*\**p* < 0.01, \*\*\**p* < 0.001, \*\*\*\**p* < 0.0001.

To identify key genes associated with chronological age, we performed a more detailed analyses of the brown and darkturquoise modules. First, a total of 924 differentially expressed genes (DEGs) in the 19 Chinese PBMC transcriptomic data were found to be dysregulated in old individuals (|logFC| ≥ 1 and adjusted *p* < 0.05). Then, as shown in Venn diagram Figure S3, 278 and 19 overlapping genes were extracted from the brown module and darkturquoise module in 19 Chinese PBMC DEG dataset, respectively. Subsequently, the protein protein interaction (PPI) network among the overlapping genes was established to identify potential aging related hub genes by using the STRING database. And then, based on the Maximal Clique Centrality (MCC) scores calculated by cytoscape, the top six highest-scored genes, including probable ATP-dependent RNA helicase (DDX27), Signal recognition particle subunit (SRP68), E3 ubiquitin-protein ligase (RNF25), Transmembrane protein 131-like (TMEM131L), UDP-N-acetylglucosamine--peptide N-acetylglucosaminyltransferase 110 kDa subunit (OGT), Early endosome antigen 1 (EEA1), exhibiting the highest connections with other genes were identified for further investigation (Figure 3D, E). Strikingly, the mRNA abundance of these hub genes was significantly associated with chronological age (Figure 3F, G). It was previously demonstrated TMEM131L could regulate immature single-positive thymocyte proliferation arrest by acting through mixed Wnt-dependent and -independent mechanisms [8]. Reports also demonstrated O-GlcNAc transferase (OGT) level was decreased in multiple aged tissues and suggested that dysregulation of OGT related O-GlcNAc formation might play an important role in the development of age-related diseases [9]. Researchers also reported the abundance of EEA1 proteins was altered in the brains of aged mice [10]. Moreover, SRP68 has been reported for its association with cellular senescence, while the ubiquitination-related genes RNF25 is not clear in immune aging. These data supported the notion that TMEM131L, OGT, EEA1, DDX27, SRP68 and RNF25 played important roles during PBMC aging, which might function as the novel candidate biomarkers of aging for Chinese individuals.

### Age-related transcriptional variation of Caucasians

Similarly, to investigate the aging-related gene modules in PBMC transcriptomes in Caucasian individuals, we performed WGCNA on microarray data from 113 Caucasian individuals, including 48 young (<40 years), 25 mid-age (40–65 years), and 41 health elderly (65–90 years). Then, a total of 16,376 genes from these transcriptomic data were used for this computation. Twenty major gene modules (Each module containing ≥160 genes) were identified. Similarly, we plotted the heatmap of module-trait relationships to evaluate the association between each module and the trait of age and sex (Figure 4A; Supplementary Table 9). The results revealed that the brown and turquoise module were found to have the highest association with chronological age (brown module: r = 0.52, *p* = 2.45e-09; turquoise module: r = −0.47, *p* = 2.05e-07). More interestingly, these two aging-related modules showed two periods in the human lifespan during which the immune system underwent abrupt changes: (i) a timepoint in early thirties, and (ii) a later timepoint after 65 years of age (Figure 4B, C). GO functional enrichment analyses suggested that the brown and turquoise modules were mainly enriched in hormone transport and postsynaptic specialization, respectively (Figure 4D, Supplementary table 10). Moreover, KEGG pathway enrichment analyses showed that the genes of brown module were mainly categorized into long-term depression and gap junction, while the turquoise module was mainly enriched in phototransduction and hedgehog signaling pathway (Figure 4E). Next, we focused on the core genes of the brown and turquoise modules. By using the differential expression analyses, we identified 1185 genes differentially expressed with chronological age in Caucasian, and 50 and 177 of these DEG genes were members of the brown and turquoise module, respectively (Figure 4F). Subsequently, the 50 and 177 genes from brown and turquoise module were subject to hub gene identification using the STRING database, respectively (Figure 4G). The results showed that the top two hub genes (Adenylate Cyclase 4, ADCY4; Phosphatidylinositol 4,5-bisphosphate 3-kinase catalytic subunit alpha isoform, PIK3CA) in turquoise module were significantly down-regulated in PBMCs of old adults (Figure 4H), whereas immunoglobulin superfamily DCC subclass member 2 (NEO1) from brown module showed the opposite result in the Caucasian cohorts (Figure 4H). From the aging atlas website (https://bigd.big.ac.cn/aging/age_related_genes), ATP Pyrophosphate-Lyase 4 (ADCY4) and Serine/Threonine Protein Kinase (PIK3CA) have both involved in Longevity regulating pathway. As reported, ADCY4 catalyzed the formation of the signaling molecule cAMP in response to G-protein signaling [11], and PIK3CA participated in cellular signaling in response to various growth factors, which also involved in the activation of AKT1 upon stimulation by receptor tyrosine kinases ligands such as EGF, insulin, IGF1, VEGFA and PDGF [12]. Besides, Gulati *et al.* reported neogenin-1 (NEO1) was associated with the long-term HSCs (LT-HSCs) expand during age [13]. Taken together, these data also revealed that ADCY4, PIK3CA, and NEO1 were critical in aging, which might serve as the novel candidate biomarkers in Caucasian individuals during the PBMC aging.

**Figure 4.**
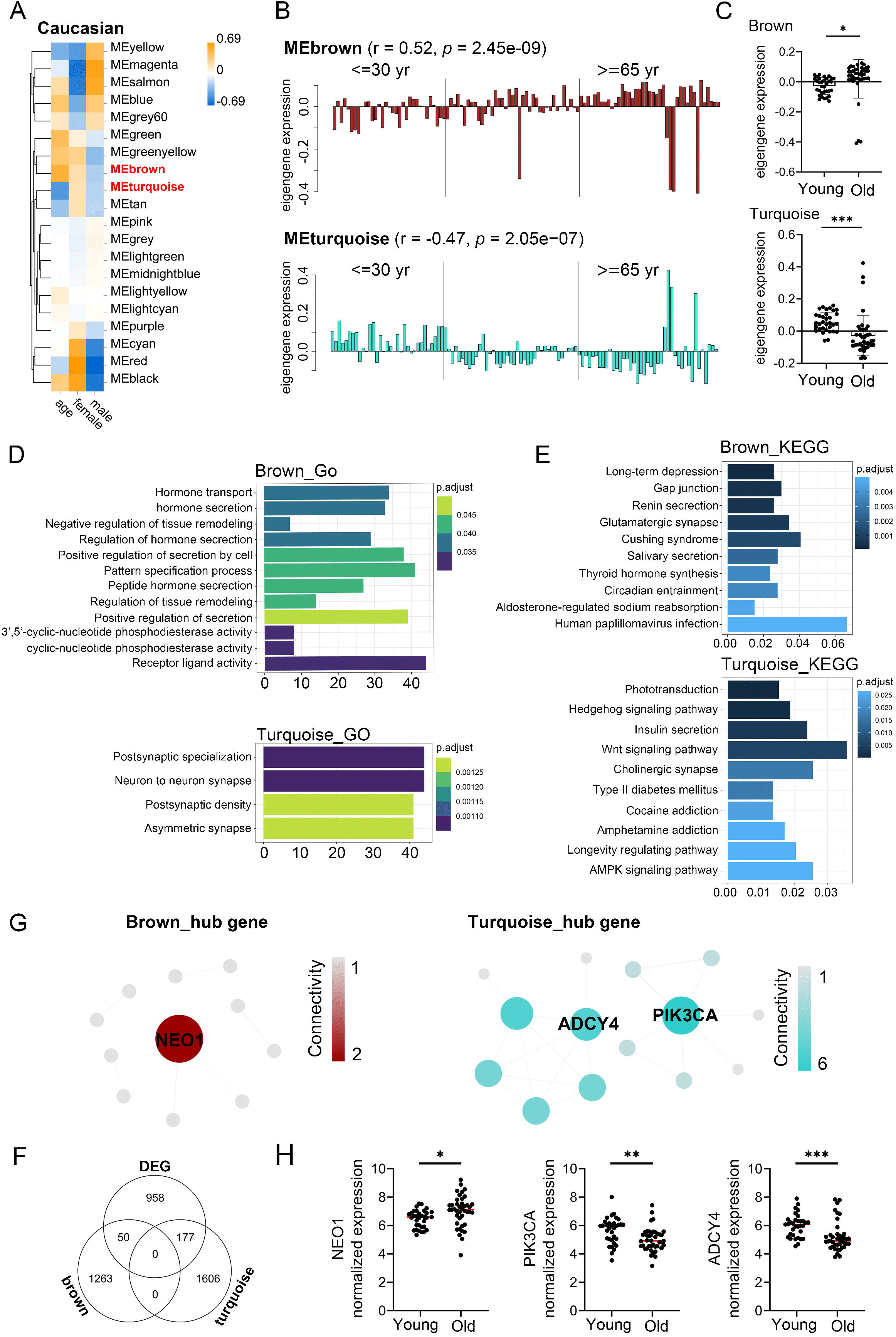
Aging-related changes in PBMC transcriptomes in Caucasian and its hub gene detection. Transcriptome data of 113 healthy Caucasian subjects in 10KIP were analyzed and gene modules were constructed by WGCNA. (A) Module-trait relationships. Row and column corresponded to module eigengenes and clinical trait (age and gender), respectively. Each cell contained the corresponding correlation and *p*-value. (B) The histograms presentation of the eigengene expression in the brown and turquoise modules from young to old. (C) Comparison of eigengene expression of the brown and turquoise modules between young and older. The transcriptomic expression of age related modules changed significantly among young and old individuals. The *p*-value was calculated by the student’s t-test, n=113, \**p* < 0.05, \*\**p* < 0.01, \*\*\**p* < 0.001. (D-E) Gene Ontology (GO) and KEGG enrichment analyses for the genes in the brown and turquoise modules. The top 10 of the GO enriched biological process and enriched KEGG pathway were shown. The color represented the adjusted *p*-values, and the size of the bars represented the gene number. (F) The Venn diagram of genes among differential expression genes (DEG) lists and co-expression module. In total, 50 and 177 overlapping genes were listed in the intersection of DEG lists and two co-expression modules. (G) Hub genes detection for the brown and turquoise modules by using the STRING database. (H) The verification of the hub genes. The top of genes in brown and turquoise modules were selected and mRNA abundance of these hub genes were detected in young and old individuals. The *p*-value was calculated by the student’s t-test, young=33, old=41, \**p* < 0.05, \*\**p* < 0.01, \*\*\**p* < 0.001, \*\*\*\**p* < 0.0001.

### Shared transcriptomic signatures of aging between Asian (Chinese) and Caucasian

As age-expectation is ethnicity dependent [14], we sought to test whether gene expression in PBMC of aging individuals differed across racial/ethnic groups. The brown module from Asian (Chinese) and the turquoise module from Caucasian were both negatively correlated with chronological age. This naturally led us to compared these two modules for common expressed genes. Ninety-five genes were shared between Asian (Chinese) and Caucasian, despite thousands of race-specific genes associated with aging (2623 and 1688 genes in Asian and Caucasian, respectively; Figure 5A). Functional annotation of these 95 shared genes revealed that they were highly enriched in the GO biological process of hindbrain development and coenzymeA metabolic process, as well as in the KEGG pathway of TGF-beta signaling (Figure 5B, C). To uncover potential regulators of common transcriptomic variation in Asian (Chinese) and Caucasian, we identified hub genes by using the STRING database. Accordingly, the top-scored genes, including peroxisomal coenzyme A diphosphatase (NUDT7) and caseinolytic peptidase B protein homolog (CLPB) were selected as the hub genes (Figure 5D). Meanwhile, two genes (OXNAD1 and MLLT3) that shared between Caucasian and Asian (Chinese) aging-related modules showed common differential expression (Figure 5E). These two potential aging-specific markers (OXNAD1 and MLLT3) were both downregulated in old Asian (Chinese) and Caucasian (Figure 5F). These data uncovered that despite the stark contrast between races in aging-related gene expression pattern, our analyses were able to highlight shared aging biomarkers with common functional enrichment in Asian (Chinese) and Caucasian.

**Figure 5.**
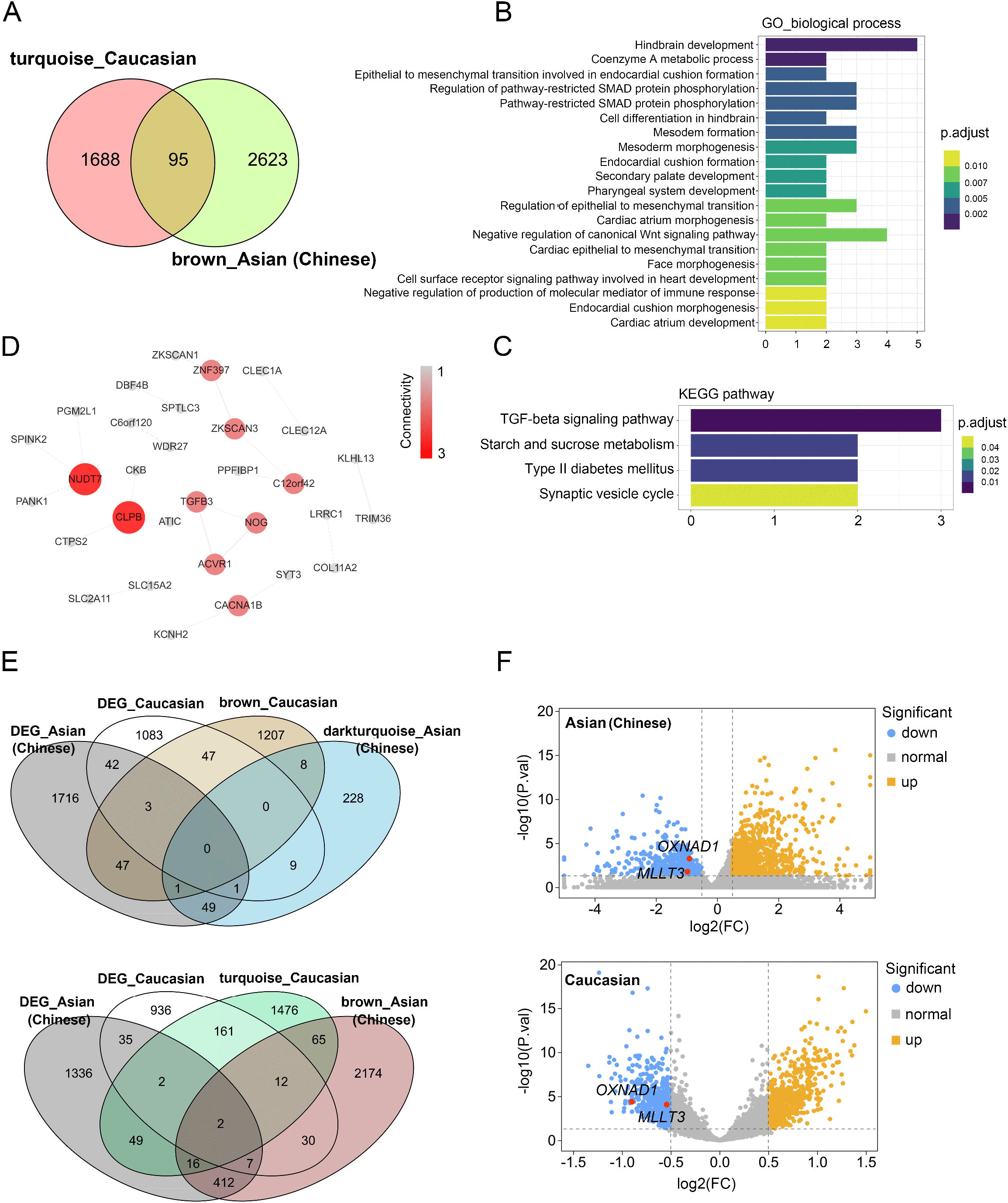
Shared transcriptomic signatures of PBMC aging between Caucasian and Asian (Chinese). The brown module from Asian (Chinese), the turquoise module from Caucasian and its differentially expressed genes (DEGs) were compared and listed. (A) The Venn diagram of genes among turquoise module from Caucasian and brown module from Asian (Chinese). Despite thousands of race-specific gene associated with aging corresponding to 2623 and 1688 genes in Asian (Chinese) and Caucasian, 95 genes in Asian (Chinese) and Caucasian significantly overlapped. (B-C) GO and KEGG enrichment analyses for the 95 common shared genes. (D) Hub genes detection for 95 genes by using the STRING database, and visualized by the cytoscape. (E) The Venn diagram of differentially expressed genes (DEGs) with the age-related modules in White and Caucasian revealed two aging-specific gene markers. (F) The two overlapping genes (OXNAD1 and MLLT3) were both downregulated, as shown in the volcano diagram for the DEG genes in Asian and Caucasian dataset.

### Validated shared genes involved in PBMC aging

After the 4 hub genes (NUDT7, CLPB, OXNAD1, MLLT3) together shared in Asian (Chinese) and Caucasian, we verified the expression levels of the hub genes among the individuals using the RNA-seq data and qPCR assay. All of the 4 hub genes were found to be significantly downregulated in old individuals compared with the youth in Asian (Chinese) and Caucasian (Figure 6A, B). Interestingly, they were all down-regulated in women during their lifespan in both Caucasian and Asian (Chinese) (Figure 6C). To further investigate whether these 4 hub genes expressed differentially during PBMC aging, we measured mRNA levels of these four hub genes (NUDT7, CLPB, OXNAD1 and MLLT3) in extracts of PBMC from 7 young adult (ages 21~30), and 5 aged health adults (ages 74+). Similarly, the mRNA level of NUDT7, CLPB, OXNAD1 and MLLT3 were both remarkably down-regulated in the elderlies (Figure 6D). All the above-mentioned observations confirmed down-expression of NUDT7, CLPB, OXNAD1 and MLLT3 is associated with PBMC aging in Asian (Chinese) and Caucasian.

**Figure 6.**
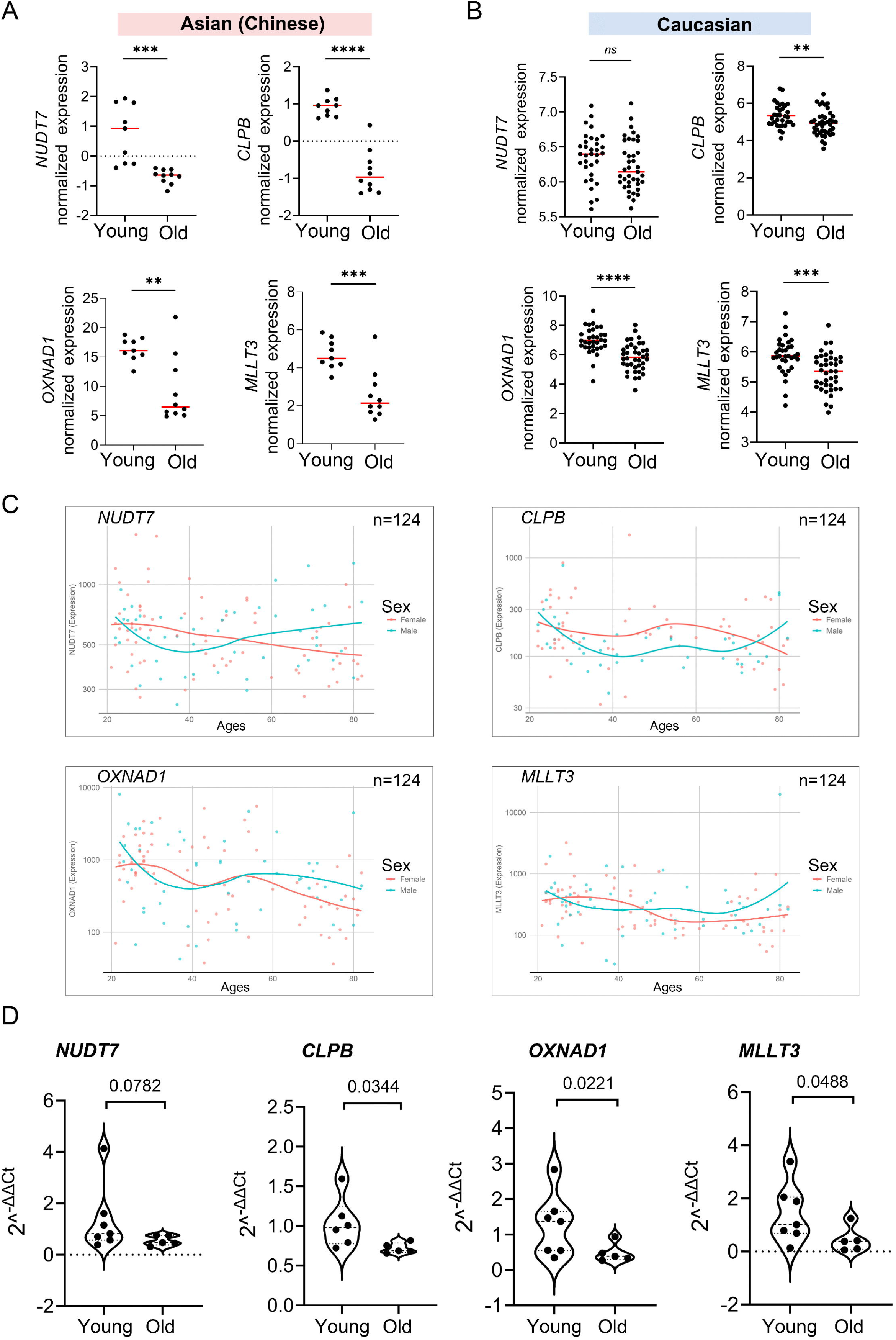
Validation of expression levels of the common hub genes involved in PBMC aging. The validation in Asian (Chinese) and Caucasian were performed using additional samples and 10KIP data, respectively. (A, B) Gene expression value of the hub genes among young and old samples in 19 Asian (Chinese) (A) and 153 Caucasian (B). Student’s t-test was used for statistical analyses. In Asian (Chinese), young n=9, old n=10; in Caucasian, young n=33, old n=40, \**p* < 0.05, \*\**p* < 0.01, \*\*\**p* < 0.001, \*\*\*\**p* < 0.0001. (C) Gene expression of hub genes among samples of man and woman during their lifespan, n=124, including 19 Asian and 105 caucasian. (D) Quantification of the four hub genes was confirmed and presented by the qPCR assay. The *p*-value was calculated by the student’s t-test. Young n=7; old n=5, \**p* < 0.05, \*\**p* < 0.01.

## MATERIALS AND METHODS

### Ethics

The study was conducted following approval by the Ethics Committee of the First affiliated Hospital of Jinan University (Approval#:KY-2020-027). All study participants provided written informed consent at baseline using institutional review board approved forms. Our study complies with Tier 1 characteristics for “Biospecimen reporting for improved study quality” (BRISQ) guidelines.

### Human subjects

All studies were conducted following approval by the Ethics Committee of Jinan University (Approval #:KY-2020-027). Following informed consent, blood samples were obtained from 31 healthy volunteers residing in the Guangzhou, China region recruited by the First affiliated Hospital of Jinan University and Guangzhou First People’s Hospital. For older adults 65 years and older, recruitment criteria were selected to identify individuals who are experiencing “usual healthy” aging and were thus representative of the average or typical normal health status of the local population within the corresponding age groups. Selecting this type of cohort was in keeping with the 2019 NIH Policy on Inclusion Across the Lifespan (NOT-98-024) [15], increasing the generalizability of our studies and the likelihood that these findings can be translated to the general population. Subjects were carefully screened in order to exclude potentially confounding diseases and medications, as well as frailty. Individuals who reported chronic or recent (i.e., within two weeks) infections were also excluded. Subjects were deemed ineligible if they reported a history of diseases such as congestive heart failure, ischemic heart disease, myocarditis, congenital abnormalities, Paget’s disease, kidney disease, diabetes requiring insulin, chronic obstructive lung disease, emphysema, and asthma. Subjects were also excluded if undergoing active cancer treatment, prednisone above 10 mg day, other immunosuppressive drugs, any medications for rheumatoid arthritis other than NSAIDs or if they had received antibiotics in the previous 6 months. Finally, smoking history data are not typically collected in these studies—including 19 Chinese individuals—since smoking is a rare habit among older adults.

### RNA-seq library generation and processing

Total RNA was isolated from PBMCs using the TRIzol (Invitrogen, United States) following manufacturer’s protocols. During RNA isolation, DNase I treatment was additionally performed using the RNase-free DNase set (Qiagen). RNA quality was checked using an Agilent 2100 Bioanalyzer instrument, together with the 2100 Expert software and Bioanalyzer RNA 6000 pico assay (Agilent Technologies). RNA quality was reported as a score from 1 to 10, samples falling below threshold of 8.0 being omitted from the study. cDNA libraries were prepared using the TruSeq Stranded Total RNA LT Sample Prep Kit with Ribo-Zero Gold (Illumina) according to the manufacturer’s instructions using 100 ng or 500 ng of total RNA. Final libraries were analyzed on a Bioanalyzer DNA 1000 chip (Agilent Technologies). Paired-end sequencing (2 × 150 bp) of stranded total RNA libraries was carried out in Illumina NovaSeq 6000. Quality control (QC) of the raw sequencing data was performed using the FASTQC tool, which computes read quality using summary of per-base quality defined using the probability of an incorrect base call. According to our quality criteria, reads with more than 30% of their nucleotides with a Phred score under 30 were removed, whereas samples with more than 20% of such low-quality reads were dropped from analyses. Reads from samples that pass the quality criteria were quality-trimmed and filtered using trimmomatic. High-quality reads were then used to estimate transcript abundance using RSEM. Finally, to minimize the interference of non-messenger RNA in our data, estimate read counts were renormalized to include only protein-coding genes.

### Microarray data acquisition

The microarray-based expression from 10 KIP provided by Lu et al. [16], was downloaded from the 10,000 immunomes project (10KIP, http://10kimmunomes.org/). This dataset contained quantile normalized genome-wide expression profiles of 153 adult human PBMC samples from young to old adults, including samples from 65 young (ages 21~40: 24 men, 41 women), 40 middle-aged (ages 41~64: 18 men, 22 women), and 48 older subjects (65+: 20 men, 28 women) and containing three races including 19 Asian, 113 Caucasian and 21 Black or Africa American.

### Identification of Key Co-expression Modules Using WGCNA

WGCNA R software package was applied to identify the co-expression modules of highly correlated genes among samples and related modules to external sample traits [17]. In our study, the gene expression data profiles of microarray data and RNA-seq profile were collected seperately to consturct gene co-expression networks by using the WGCNA package. To build a scale-free network, optimum soft powers β were selected using the function pickSoftThreshold. Next, the adjacency matrix was created by the following formula: aij = |Sij|β (aij: adjacency matrix between gene i and gene j, Sij: similarity matrix which was done by Pearson correlation of all gene pairs, β: softpower value), and was transformed into a topological overlap matrix (TOM) as well as the corresponding dissimilarity (1-TOM). Afterwards, a hierarchical clustering dendrogram of the 1-TOM matrix was constructed to classify the similar gene expressions into different gene co-expression modules. To further identify functional modules in a co-expression network, the module-trait associations between modules, and clinical trait information were calculated according to the previous study [18].

Therefore, modules with high correlation coefficient were considered candidates relevant to clinical traits, and were selected for subsequent analyses. A more detailed description of the WGCNA method was reported in our previous study [18].

### Differential Expression analyses and Interaction with the Modules of Interest

The R package limma (linear models for microarray data) provides an integrated solution for differential expression analyses on RNA-Sequencing and microarray data[19]. In order to find the differentially expressed genes (DEGs) between Asian and Caucasian, limma was applied in the Asian RNA-seq and Caucasian dataset to screen out DEGs, respectively. The p-value was adjusted by the Benjamini–Hochberg method to control the false discovery Rate (FDR). Genes with the cut-off criteria of |logFC| ≥ 0.50 and adjusted *p*< 0.05 were regarded as DEGs. The DEGs of the Asian and Caucasian dataset were visualized by a volcano plot using the R package ggplot2 (Wickham H, 2016). Subsequently, the overlapping genes between DEGs and co-expression modules, listed by the Venn diagram using the R package VennDiagram, were used to identify potential prognostic genes[20].

### Functional Annotation for the Modules of Interest

For genes in each module, Gene Ontology (GO) and KEGG pathway enrichment analyses were conducted to analyze the biological functions of modules. Significantly enriched GO terms and pathways in genes in a module comparing to the background were defined by hypergeometric test and with a threshold of false discovery rate (FDR) less than 0.05. The *clusterProfiler* package offered a gene classification method, namely groupGO, to classify genes based on their projection at a specific level of the GO corpus, and provided functions, *enrichGO* and *enrichKEGG,* to calculate enrichment test for GO terms and KEGG pathways based on hypergeometric distribution [6]. Thus, we input the interesting modules into the *clusterProfiler* by comparing them to the annotated gene sets libraries, with a cut-off criterion of adjusted *p* < 0.05. GO annotation that contained the three sub-ontologies—biological process (BP), cellular component (CC), and molecular function (MF)—could identify the biological properties of genes and gene sets for all organisms [6].

### Construction of PPI and Screening of Hub Genes

In our study, we used the STRING (Search Tool for the Retrieval of Interacting Genes) online tool, which was designed for predicting protein-protein interactions (PPI), to construct a PPI network of selected genes [21]. Using the STRING database, genes with a score ≥ 0.4 were chosen to build a network model visualized by Cytoscape (v3.7.2). In a co-expression network, Maximal Clique Centrality (MCC) algorithm was reported to be the most effective method of finding hub nodes [22]. The MCC of each node was calculated by CytoHubba, a plugin in Cytoscape [22]. In this study, the genes with the top 10 MCC values were considered as hub genes.

### Verification of the Hub Genes

In order to confirm the reliability of the hub genes, we tested the expression patterns of the hub genes from healthy individuals including 7 young (ages: 23~30) and 5 old (ages: □74). The expression level of each hub gene between young and old individuals was plotted as a violin graph. Total RNA from PBMCs was extracted by TRIzol (Invitrogen, United States). Synthesis of cDNA was performed by using 2 μg of total RNA with PrimeScriptTM Reverse Transcriptase (Takara) according to the manufacturer’s instructions. Specific primers used for qPCR were listed in the supplementary table 11. The gel image was acquired in the Gel Doc 1000 system and analyzed using the Quantity One software (Bio-Rad Laboratories, Hercules, CA, United States). ACTB was chosen as the endogenous control and cycle dependence was carried out to ensure that the PCR products fell within the linear range. Quantitative real-time PCR was performed using the SYBR^®^ Premix Ex Taq Kit (Takara) in a 7900 Real Time PCR System (Applied Biosystems, United States) for at least three independent experiments. The relative quantification expression of each gene was normalized to ACTB, and calculated using the 2-^ΔΔΔCT^ method.

### Statistical analyses

All statistical analyses was performed using GraphPad Prism 6 Software (GraphPad Software, San Diego, CA, United States). The data were expressed as the mean ± standard deviation (SD). Comparison between two groups was conducted by using Student’s t-test. *P*-values less than 0.05 were considered as statistically significant.

## DISCUSSION

Age-associated changes in gene expression levels point towards altered activity in defined age-related molecular pathways that might play vital roles in the mechanisms of increased susceptibility to ageing diseases. In contrast to earlier studies of human age-related molecular differences[23], we detected 184 adult individuals from all ages among Asian, Caucasian and Africa and American ancestry. In our study, a total of four significant gene modules with the same expression trends were identified using integrated bioinformatic analyses in Asian and Caucasian populations. As suggested in functional annotation analyses by the *clusterProfiler* package, these module genes were mainly enriched in virus infection, amino acid metabolic and differentiation, which were basic processes in aging mechanisms including dysregulation of herpes simplex virus 1 infection, valine, leucine and isoleucine degradation, long-term depression, gap junction, and hedgehog signaling pathway. Furthermore, according to MCC scores from the *CytoHubba plugin* in Cytoscape, the top chronological age-related genes were screened out (namely TMEM131L, OGT, EEA1, DDX27, SRP68 and RNF25 in Asian; ADCY4, PIK3CA and NEO1 in Caucasian). According to reports in the literature, all of these genes were more or less closely associated with aging. Consistent with these reports, the expression of these genes was also found be significantly regulated among young and old individuals in our study, supporting these genes might play a causal role in human PBMC aging. More importantly, the breakpoint analyses uncovered that although aging related transcriptomic changes accumulated gradually throughout adult life, there were two periods in the human lifespan during which the immune system underwent abrupt changes. The two breakpoints (30 and 65~70 years old) were much similar in Asian and Caucasian during their whole lifespan. The differences in the timing of age-related changes could be helpful in clinical decisions regarding when to start interventions/therapies.

Despite well-characterized race differences in immune responses, disease susceptibility, and lifespan, it was unclear whether aging differentially affected peripheral blood cells of European and Asian ancestry. To fill this gap, we generated RNA-seq data in PBMCs from 19 age-matched healthy adults in Guangdong province of China and downloaded microarray data of 153 individuals’ PBMCs from 10 KIP (http://10kimmunomes.org/) including European, Asian and the other ancestry groups (19 Asian, 113 Caucasian and 21 Black or Africa American). Weighted gene correlation network analyses (WGCNA) was an integrated bioinformatic analyses, which could provide a comprehensive characterization of the transcriptomic changes for disease’s functional interpretation and led to new insights into the molecular aspects of clinical-pathological factors [17]. So by using this integrated bioinformatic analyses, we discovered a genomic signature of aging that was shared between Asian (Chinese) and Caucasian ancestry including (1) 95 age-associated genes shared in 132 individuals. (2) four hub genes (NUDT7, CLPB, OXNAD1 and MLLT3) all decreased in old ages. According to reports in the literature about these four hub genes, NUDT7, acted as a coenzyme A (CoA) diphosphatase, which mediated the cleavage of CoA. NUDT7 functioned as a house-keeping enzyme by eliminating potentially toxic nucleotide metabolites, such as oxidized CoA from β-oxidation in the peroxisome, as well as nucleotide diphosphate derivatives, including NAD+, NADH, and ADP-ribose [24]. Furthermore downregulation of NUDT7 in mice accelerating senescence [25], was observed in the liver of starved mice [26]. Interestingly, OXNAD1 also known as oxidoreductase NAD-binding domain-containing protein, had been reported differentially expressed with chronological age [23]. And according to the uniprot anotation for CLPB, it might function as a regulatory ATPase and be related to secretion/protein trafficking process, involved in mitochondrial-mediated antiviral innate immunity, and activated RIG-I-mediated signal transduction and production of IFNB1 and proinflammatory cytokine IL6 [27]. Moreover, the hub gene of MLLT3 was a component of the superelongation complex and co-operated with DOT1L, which di/trimethylates H3K79 to promoted transcription [28, 29]. Recently, Vincenzo Calvanese, et.al, found MLLT3 could govern human haematopoietic stem-cell self-renewal and engraftment [30]. From above, NUDT7 and OXNAD1 both had an important role in cellular metabolism and aging, which was consistent with our finding of PBMC aging analyses, while the role of CLPB and MLLT3 in immune aging or senescence was unclear. Thus, by using co-expression networks, we identified new genes that were likely important in PBMC aging in Asian and Caucasian ancestry, opening new avenues of enquiry for future studies.

By WGCNA analyses, aging-specific regulatory modules and hub genes were identified in bulk PBMCs in different races. Although this approach was effective in annotating the aging signatures, it was prone to biases in the differences of data quality and formats. Besides, as we had much smaller sample sizes for both PBMCs in Asian and the other ancestry groups, we used a nominal *P*-value threshold (*p*<0.05) in these specific sub-analyses. Larger sample sizes will ultimately be needed to fully understand the transferability of the aging-transcriptome signatures. And more importantly, further studies were needed to verify the important molecules, identified here (NUDT7, CLPB, OXNAD1 and MLLT3) as aging specific biomarkers of immune system aging. Future studies might be needed to describe these race differences at single-cell resolution and in sorted cells and to establish their functional implications. Taken together, these findings indicated that aging played a critical role in human immune system aging and should be taken into consideration while searching for molecular targets and time frames for interventions/therapies to target aging and age-related diseases.

## Acknowledgement

This work was supported by grants from the National Key Research and Development Program of China (No.2018YFC2002003), the Natural Science Foundation of China (U1801285, 81971301), Guangzhou Planned Project of Science and Technology (201904010111, 202002020039), Milstein Medical Asian American Partnership (MMAAP) Foundation Irma and Paul Milstein Program for Senior Health Research Project Award (to G.C.), and the China Post-doctoral Science Foundation (No.55350399) and the Postdoctoral Start-up Fund from the First Affiliated Hospital of Jinan University (No.809026). We would like to thank all donors as well as the Guangzhou First People’s Hospital for provision of the samples. We thank the Beijing Novogene Biotechnology Co., Ltd. for assistance with RNA-sequencing.

## Author Contributions

The concept of the study was planned by G.C., O.J.L. and Y.H. Experiments were conducted, analyzed, and interpreted by Y.H., Y.X., L.M., W.L., G.Z., Y.H.(Yutian Hu), H.N., F.G., L.H., and L.G. Sample preparation for mRNA, and sequencing were done by Y.X., J.X. and W.L. Y.H. drafted the manuscript. O.J.L. and G.C. edited the manuscript and provided advice. All authors contributed to the article and approved the submitted version.

## Data Availability Statement

All data relevant were contained within the article. The sequencing data presented in the study has been deposited in the Sequence Read Archive (SRA) database, which was hosted by the NCBI, under accession number (SRA: PRJNA703752).

## Ethics Statement

The studies involving human participants were reviewed and approved by the local ethics committee of the First Affiliated Hospital of Jinan University. The patients/participants provided their written informed consent to participate in this study.

## Supplementary information

### Supplementary Figures

Supplementary Figure S1. The age and race distribution of 153 individuals from 10 KIP dataset.Supplementary Tables.

Supplementary Figure S2. Power values and module clustering of WGCNA for the transcriptome data of 153 healthy human subjects in 10KIP.

Supplementary Figure S3. The venn diagram of genes among DEG lists and co-expression module in 19 Asian (Chinese) RNA-seq data.

### Supplementary tables

Supplementary Table 1: Basic information for 19 Chinese healthy individuals.

Supplementary Table 2: Basic information for 153 individuals from 10 KIP.

Supplementary Table 3: Trait-module relationships of the 153 individuals.

Supplementary Table 4: Module eigengene (ME) of the 153 individuals.

Supplementary Table 5: PCA analysis for the transcriptome of 153 individuals.

Supplementary Table 6: Trait-module relationships for the 19 Asian (Chinese).

Supplementary Table 7: Module eigengene (ME) of the 19 Asian (Chinese).

Supplementary Table 8: GO and KEGG enrichment for brown and Darkturquoise modules from 19 Chinese.

Supplementary Table 9: Trait-module relationships in the 113 Caucasian.

Supplementary Table 10: GO and KEGG enrichment for brown and turquoise modules from 113 Caucasian.

Supplementary Table 11: The qPCR primers for homo sapiens.

