## Supplementary File for "Identification of age and ethnicity specific gene expression biomarkers for immune aging"

[Yang Hu](https://www.ncbi.nlm.nih.gov/pubmed/?term=Hu%20Y%5bAuthor%5d&cauthor=true&cauthor_uid=30210331)^1,2^, Yudai Xu^1^, Lipeng Mao^1,3^, Wen Lei^1^, Jian Xiang^1^, Guodong Zhu^4^, Yutian Hu^5^, Haitao Niu^6^, Feng Gao^7^, Lijuan Gao^1^, Li`an Huang^2^, Oscar Junhong Luo^1,3^*, Guobing Chen^1,2^*

1, Institute of Geriatric Immunology, Department of Microbiology and Immunology, School of Medicine, Jinan University, Guangzhou, Guangdong, China.

2, Department of Neurology, the First Affiliated Hospital, Jinan University, Guangzhou, Guangdong, China.

3, Department of Systems Biomedical Sciences, School of Medicine, Jinan University, Guangzhou, Guangdong, China.

4, Department of Geriatrics, Guangzhou First People's Hospital, School of Medicine, South China University of Technology, Guangzhou, Guangdong, China.

5, Meng Yi Center Limited, Macau, China

6, School of Medicine & Institute of Laboratory Animal Sciences, Jinan University, Guangzhou, Guangdong, China

7, School of Medicine, Jinan University, Guangzhou, Guangdong, China.

*Correspondence should be addressed to: Drs. Oscar Junhong Luo or Guobing Chen, Institute of Geriatric Immunology, School of Medicine, Jinan University, 601 Huangpu Avenue West, Guangzhou 510632, Guangdong Province, China..

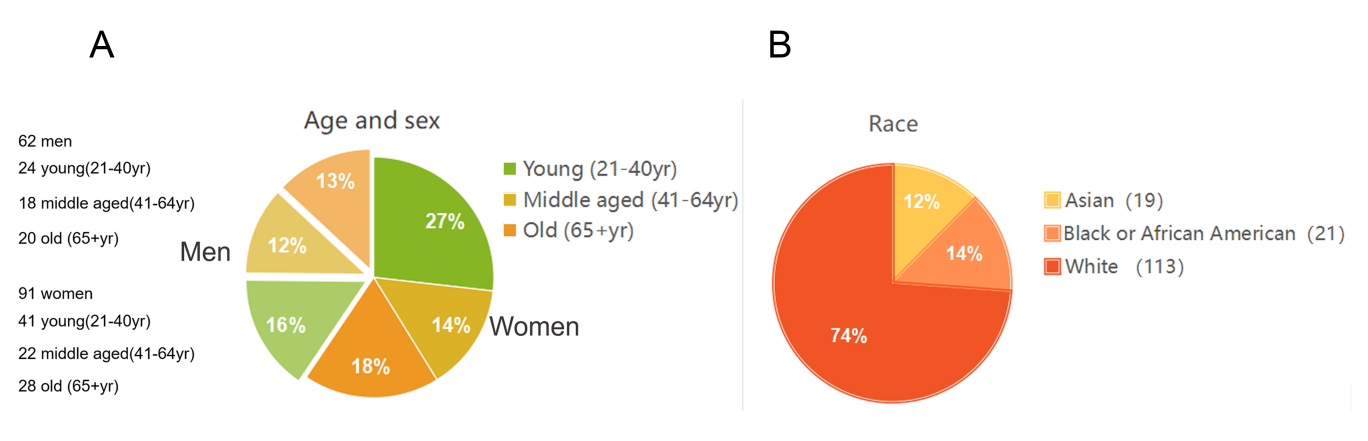

**Supplementary Figure S1. The age and race distribution of 153 individuals from 10 KIP dataset.** (A) The component of age among the 153 samples. (B) The component of race among 153 samples.

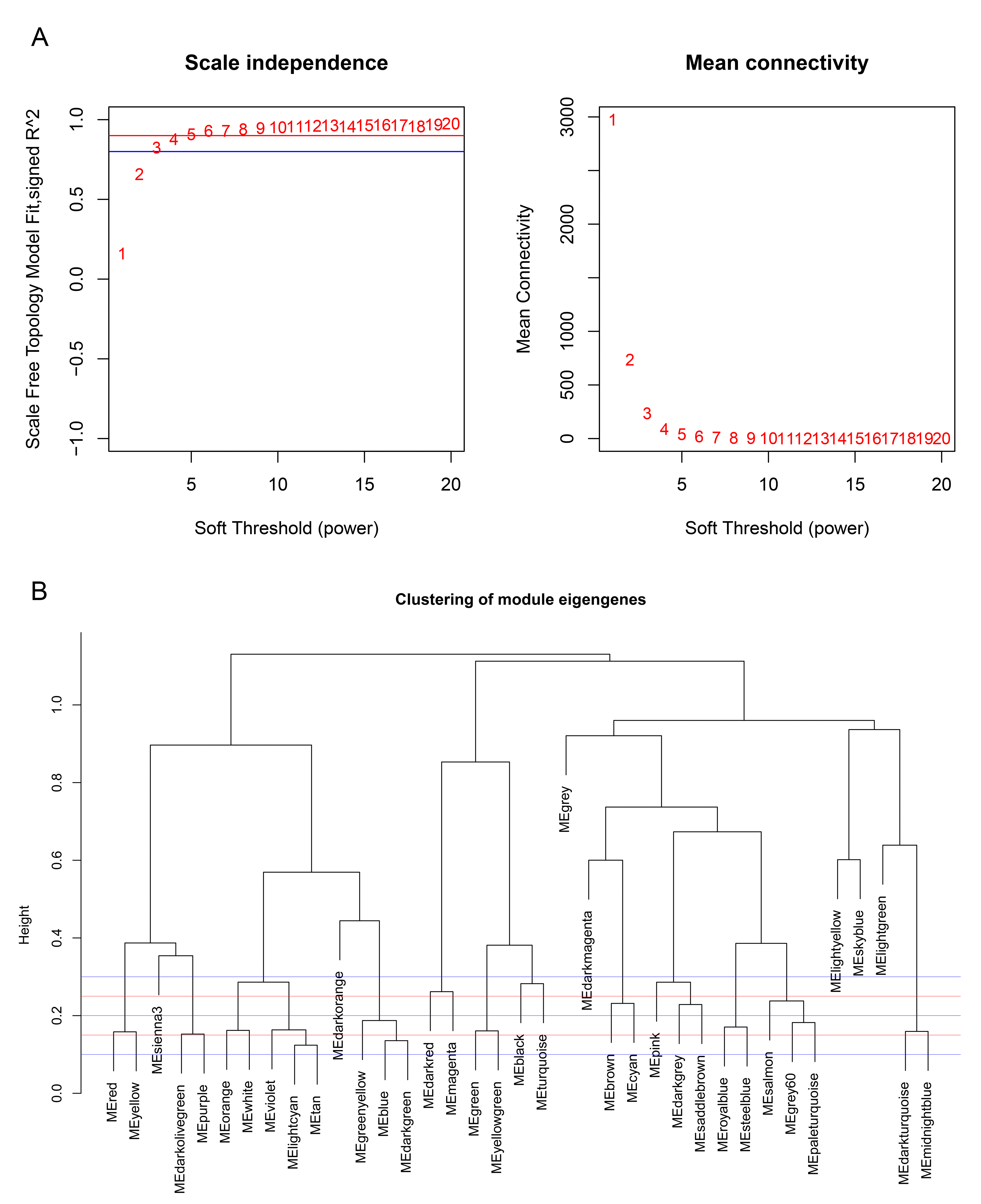

**Supplementary Figure S2. Power values and module clustering of WGCNA for the transcriptome data of 153 healthy human subjects in 10KIP.** (A) Selection of the soft-thresholding powers. The left panel showed the scale-free fit index versus soft-thresholding power. The right panel displayed the mean connectivity versus soft-thresholding power. Power 6 was chose for which the fit index curve flattens out upon reaching a high value (>0.9). (B) Meta-module identification. The module network dendrogram was constructed by clustering module eigengene distances. The horizontal line (blue and red line) represents the threshold (0.2) used for defining the meta-modules.

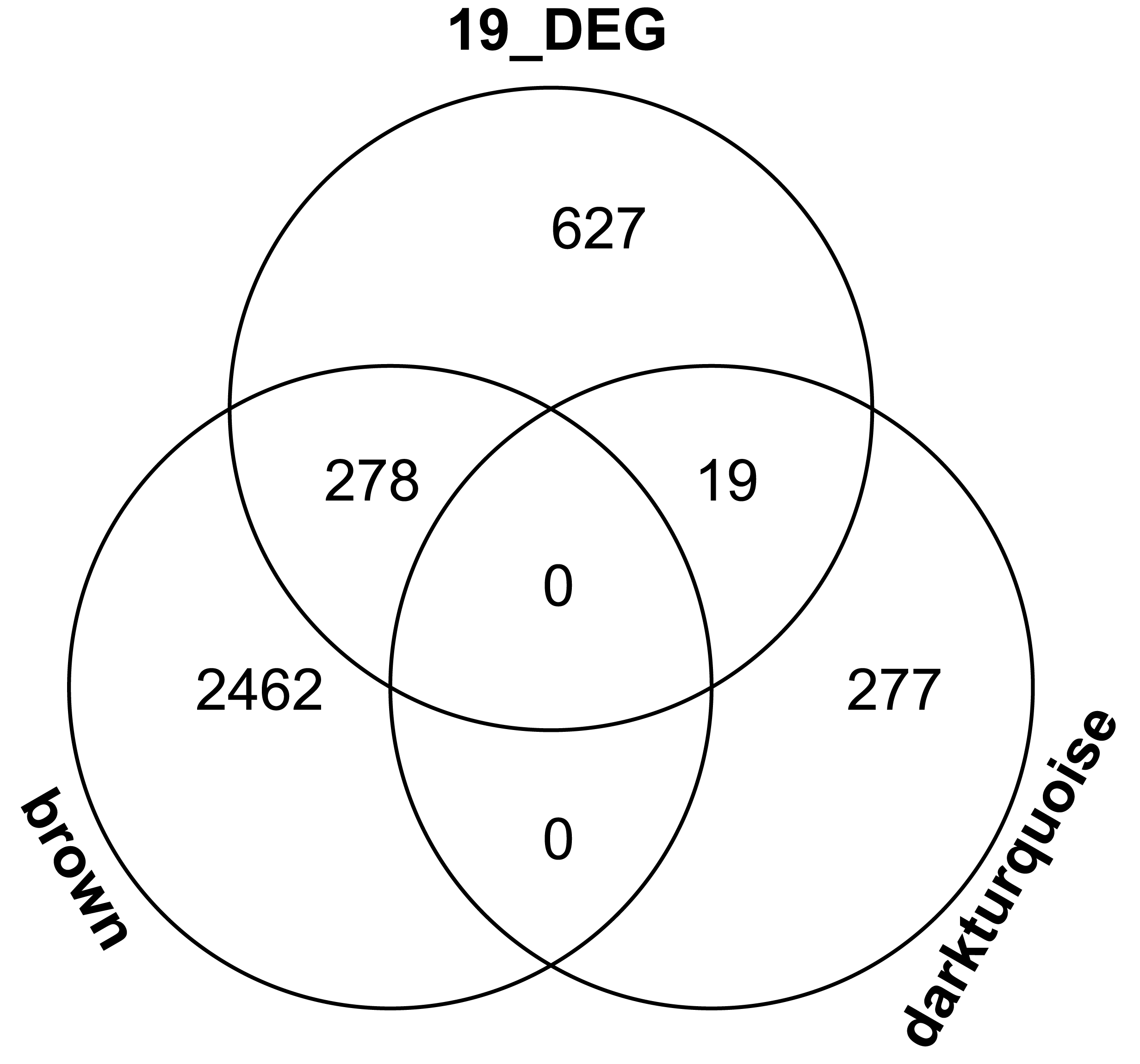

**Supplementary Figure S3. The venn diagram of genes among DEG lists and co-expression module in 19 Chinese RNA-seq data.** In total, 278 and 19 overlapping genes were listed in the intersection of DEG lists and two co-expression modules.
